# Synergies and trade-offs between maintaining climate niche variability and preserving climate stability

**DOI:** 10.1101/2025.10.26.684389

**Authors:** Marta Cimatti, Thiago Cavalcante, Heini Kujala, Sara Si-Moussi, Wilfried Thuiller, Piero Visconti, Moreno Di Marco

## Abstract

The impact of climate change on biodiversity accelerates, calling for climate resilient conservation strategies such as protecting areas of high climatic stability (i.e. climate refugia) or protecting the variability of species climatic niches (to preserve adaptive potential). Developing a spatial framework that integrates both strategies, we identify priorities to protect climatic niche components of 1,207 European vertebrates. Priority areas for protecting climatic niches under low climate velocity or low magnitude were respectively found in mountainous/southern regions and in northern/eastern Europe. These synergy areas overlapped by 48–73% with single-objective prioritizations focused on either niche components or climatic stability. Trade-offs occur where climatic niches diversity is high but climate stability is low, such as eastern Europe (velocity) or the Mediterranean and North Fennoscandia (magnitude). Our results reveal spatial mismatches between climate refugia and spatial priorities to preserve adaptive potential, emphasizing the need to combine both strategies in conservation planning.

## Introduction

As biodiversity loss and the degradation of ecosystem services gain increasing prominence on the international agenda, the European Union (EU) has adopted a comprehensive biodiversity strategy centred on area-based conservation ^1,2^. A key objective of this strategy is to protect 30% of terrestrial and marine areas, with one-third under strict protection, through a coherent and climate-resilient Trans-European Nature Network (TEN-N). EU guidance emphasizes the importance of protecting and connecting ecosystems in ways that build resilience to climate change. In fact, while habitat loss has historically been the leading driver of biodiversity decline ^3^, climate change has recently emerged as an equally critical threat to species and ecosystems ^4,5^. Under shifting climatic conditions, the effectiveness of protected areas (PAs) is increasingly compromised, particularly where static boundaries fail to align with future habitat suitability ^6^.

Species may exhibit plastic responses to climate change, such as shifts in phenology, reproductive timing, or ecological behaviour ^4,7,8^. Yet, long-term persistence will depend on their capacity to adapt to the new climatic conditions through evolutionary processes or, alternatively, track favourable conditions through range shifts. When these mechanisms are insufficient, climate-driven declines may undermine conservation efforts ^9,10^. To ensure long-term conservation success, the development of reserve networks through newly established PAs must explicitly account for such climate dynamics ^6,11,12^. Various strategies have been presented for improving the climate resilience of PA networks ^13^, such as: (i) protecting climate refugia, areas expected to retain stable climatic conditions despite broader environmental change ^14^; (ii) protecting a set of areas that maximise the representation of climatic variability within each species’ range, thus maintaining the environmental heterogeneity that maintain genetic diversity and favour adaptive potential ^15^.

Climate refugia serve as “safe havens” for species, buffering them from the most severe climate impacts ^16^. Such areas often harbour high genetic diversity and unique evolutionary lineages, having supported species persistence through past climatic fluctuations ^17–19^. In parallel, maintaining habitats spanning a range of climatic conditions within species’ distributions helps safeguard intraspecific variation, which is essential for adaptation. Recent studies suggest that species harbouring broad environmental variability within their distribution are indeed better equipped to adapt to the changing conditions locally, through phenotypic and genotypic variation critical for evolutionary resilience, instead of being forced to shift their ranges ^15,20–23^. This may be advantageous particularly in human dominated areas, such as Europe, where dispersing through highly fragmented landscapes to track suitable conditions is challenging for species ^24,25^. Together, these insights suggest that a climate-resilient PA network should seek to balance two objectives: protecting areas with high predicted climatic stability (refugia), and areas that together best represent the diversity of species’ climatic niches across the landscape, hereafter called bioclimatic niche diversity. Importantly, while the first objective is area-specific (not targeted to any species in particular) the second one is species-specific (focusing on the climatic adaptation of each species). Here, we propose a generalizable spatial framework (Fig. 1) that identifies priority areas for contributing to both objectives, and we apply it to 1270 species of European tetrapods.

**Fig. 1:**
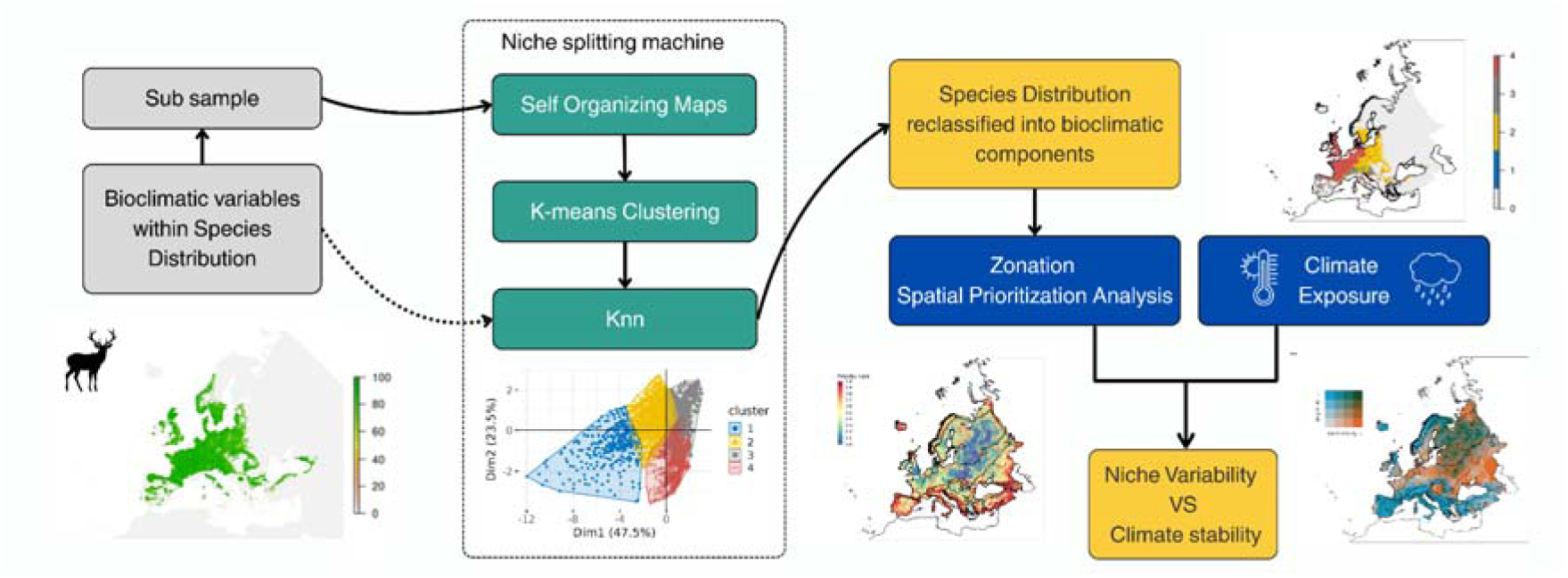
Methodological framework for selecting synergistic areas and evaluating trade-offs between species bioclimatic niche variability and climatic stability in terms of climate velocity and magnitude.

Building on modelled species distributions, we identify distinct bioclimatic components within each species’ range. The components are then used as conservation features in a spatial prioritization to identify areas that together best represent the diversity of species’ climatic niches across the landscape, with the goal of maximising species’ adaptive capacity under climate change. The high-priority conservation areas are defined as the top 30% ranked cells that maximize the protection of bioclimatic components across species. We then intersect the priority areas with future climate stability (both in terms of low velocity and magnitude) to evaluate synergies and trade-offs between maintaining bioclimatic niche variability and prioritizing climate stability.

## Results

### Identification of bioclimatic components

We have delineated the bioclimatic components of each European tetrapod species, based on their modelled distribution at 1 km resolution extracted from the EBV data portal ^26^. Starting from each species’ distribution model, we employed Self-Organizing Maps (SOMs) to identify a reduced set of local climate nodes and then applied a k-means clustering to individual components from each other. Each pixel within a predicted species’ distribution was then assigned to the most appropriate bioclimatic component using K-Nearest Neighbours (KNN). The clustering yielded a mean value of five bioclimatic components per species, and a maximum number of nine for the azure tit (*Cyanistes cyanus*) (Fig. S2.1). As a result we split 1,050 out of 1027 species (excluding species with range size smaller than 20,000 km²,^27^) into 4,281 bioclimatic components. Precipitation seasonality (bio15) and temperature seasonality (bio14) emerged as the most influential variables for delineating within-species bioclimatic components, particularly for Amphibia and Reptilia. Annual precipitation (bio12) and growing degree days (gdd5) also showed moderate to high importance across all taxa, especially for birds and mammals (Fig. S2.2). Mean classification accuracy (0.97, sd=0.013) and Cohen’s Kappa (K=0.96, sd=0.015) indicated that the clustering reliably captured the distinct bioclimatic components within species’ modelled ranges. The clustering validation metrics (silhouette coefficient, Dunn index, Calinski– Harabasz, and Davies–Bouldin) generally supported the validity of the identified clusters across species, indicating reasonable within-cluster coherence and separation ( Fig. S2.1; see also Supplementary Material).

Among mammals, the red deer (*Cervus elaphus*) is a widespread species with a predicted distribution covering 2.7 million km² of suitable habitat in Europe. Due to the large area, we subsampled 1 million grid cells before applying the SOM, which aggregated the species’ bioclimatic space into 5,000 nodes. The optimal number of clusters was four (Fig. 2), and the resulting k-means clustering produced moderately overlapping groups, with a median silhouette value of 0.33 (on a -1 to +1 range) and only 2.8% negative values, indicating an overall coherent classification. Snow cover duration was the most critical factor influencing red deer clusterization (Fig. S2.3). This reflects the species’ strong dependence on winter habitats with limited snow persistence for foraging access. The greater spotted eagle (*Clanga clanga*) also has a broad distribution, spanning approximately 2.13 million km², which was divided into 7 unique clusters (median silhouette = 0.31, 3.32% negative values). The populations of spotted eagle are mostly distinguished by temperature seasonality (bio4) suggesting population level adaptation to specific thermal regimes, followed by growing degree days (gdd5) and snow water equivalent (swe, Fig. S2.3).

**Fig. 2:**
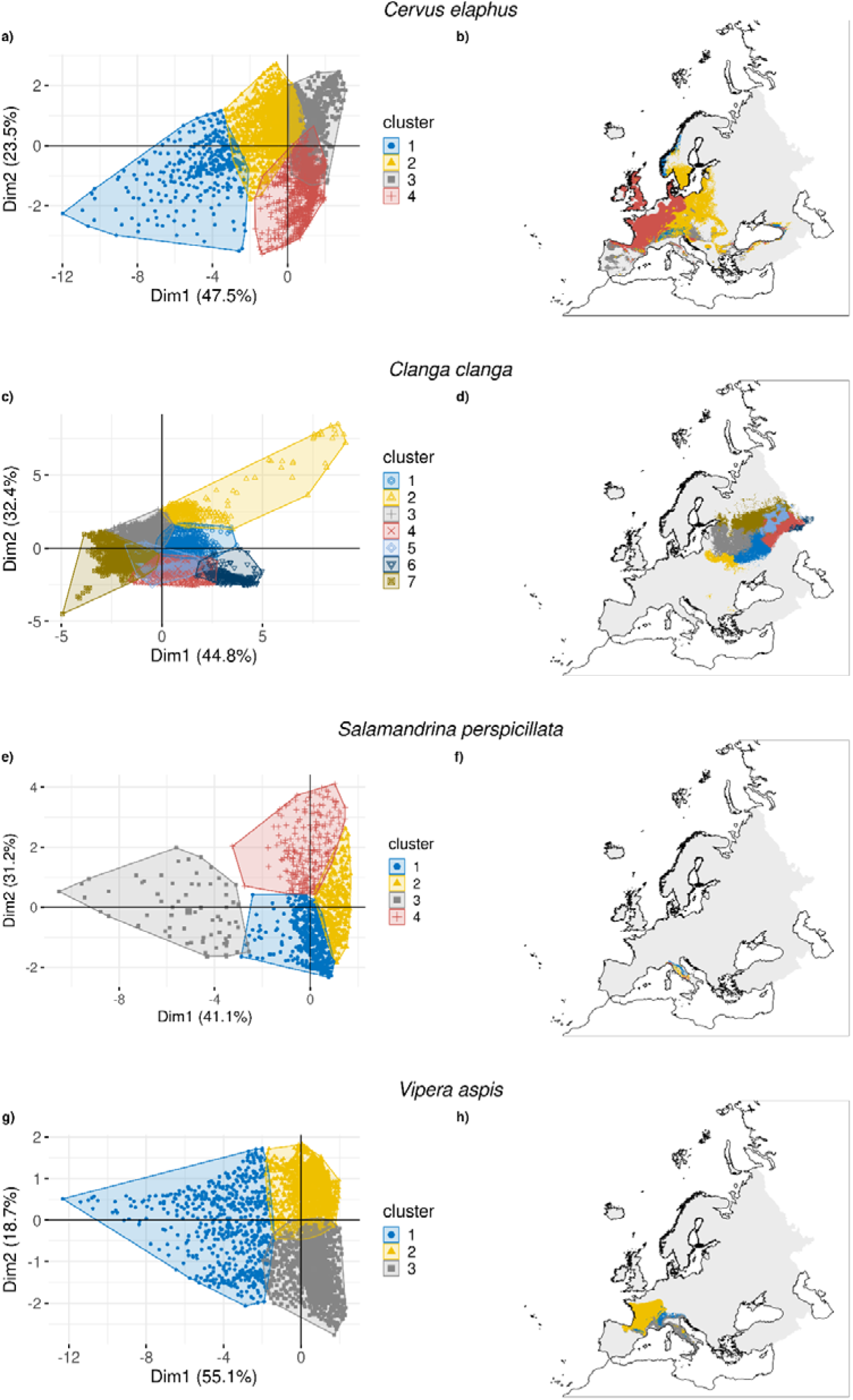
Example illustrating the distribution of sampled locations in the bioclimatic space for the range of the species on the two dominant axis of a Principal Component Analysis (PCA) a), c), e), g) and the range of the species classified into separate bioclimatic clusters b), d) f), h) for Cervus elaphus, Clanga clanga, Salamandrina perspicillata, Vipera aspis. The coloured areas represent the identified bioclimatic clusters (components), and similarly coloured symbols locations within the distribution that fall into these clusters.

In contrast, the spectacled salamander (*Salamandrina perspicillata*) has a much smaller and fragmented distribution of 51,503 km². Despite the limited range, the clustering showed a reasonably coherent structure with a median silhouette value of 0.29 and 4.51% negative values, indicating some uncertainty. Population groupings were primarily driven by variability in seasonal rainfall patterns (precipitation seasonality, bio15, with a misclassification rate of ∼0.25, Fig. S2.3) which aligns with the species’ known dependence on stable moisture regimes for reproduction and cutaneous respiration. Variability in seasonal rainfall also explains the bioclimatic clusters of the asp viper (*Vipera aspis*), followed by temperature seasonality (bio4) (misclassification rates ∼0.3 and ∼0.2, respectively; Fig. S2.3). These variables likely reflect the species’ occurrence across regions with contrasting rainfall regimes and temperature fluctuations, indicating ecological differentiation in how populations cope with moisture and thermal conditions. These drivers divide the distribution of the asp viper (687,172 km²) into three clearly separated and coherent bioclimatic clusters (median silhouette = 0.30; 2.15% negative values; Fig. 2).

### Comparison of species bioclimatic components in conservation prioritization and climate metrics

The high-priority conservation areas that best represent the diversity of species’ bioclimatic niches were primarily concentrated in mountainous regions, coastal and island areas, especially in the Mediterranean and Caspian sea (Fig. 3a). In contrast, low-priority areas were predominantly located in intensively cultivated lowland regions, particularly across Western and Central Europe. Instead, areas with both high local climate velocity and magnitude were mostly concentrated in northeastern Europe, particularly western Russia, and parts of Fennoscandia (Fig. 3b). Areas with both low climate velocity and magnitude, indicating relative climatic stability, spanned most biogeographic regions. This result reveals high potential for synergies with biodiversity priorities, making it possible to select priority areas from regions with low climate risk in most cases (i.e. areas of synergy).

**Fig. 3:**
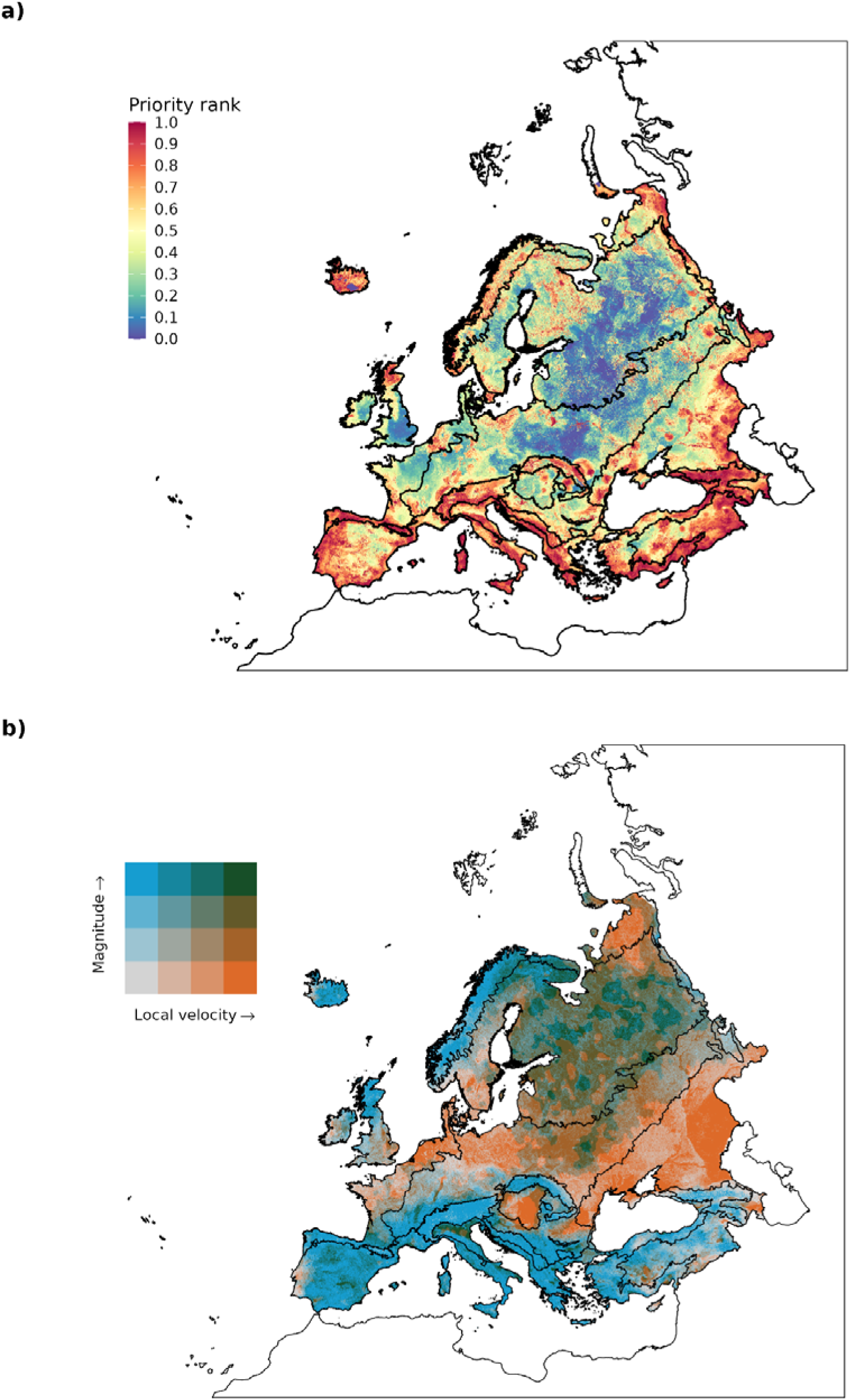
Prioritization map vs climate risk map. Panel a) presents a spatial conservation prioritization map. The prioritization ranking is represented by a colour gradient, where red areas indicate the high conservation priority and blue areas represent the low priority. Panel b)presents a bivariate representation of climate metrics across Europe under the SSP3-7.0 scenario. Dark green areas highlight regions where both local velocity and climate magnitude are high.

### Synergies and Trade-offs in Climate Resilience Conservation Planning

When looking at climate stability in terms of low climate velocity, synergy areas were mostly found in mountainous regions, such as the Alps, Carpathians, Pyrenees, the Scandinavian Mountains, parts of the Balkans, and the Caucasus (Fig. 4a). When stability was defined in terms of low climate magnitude, synergy areas shifted towards northern and eastern Europe, including Sweden, parts of Poland, the Baltic states, western Russia, and regions north of the Black Sea and along the Caspian coast (Fig. 4b). Notably, part of the southern Iberian peninsula also emerged as potential climate refugia, suggesting more stable conditions that could support long-term species persistence despite surrounding climatic instability (Fig. 4b).

**Fig. 4:**
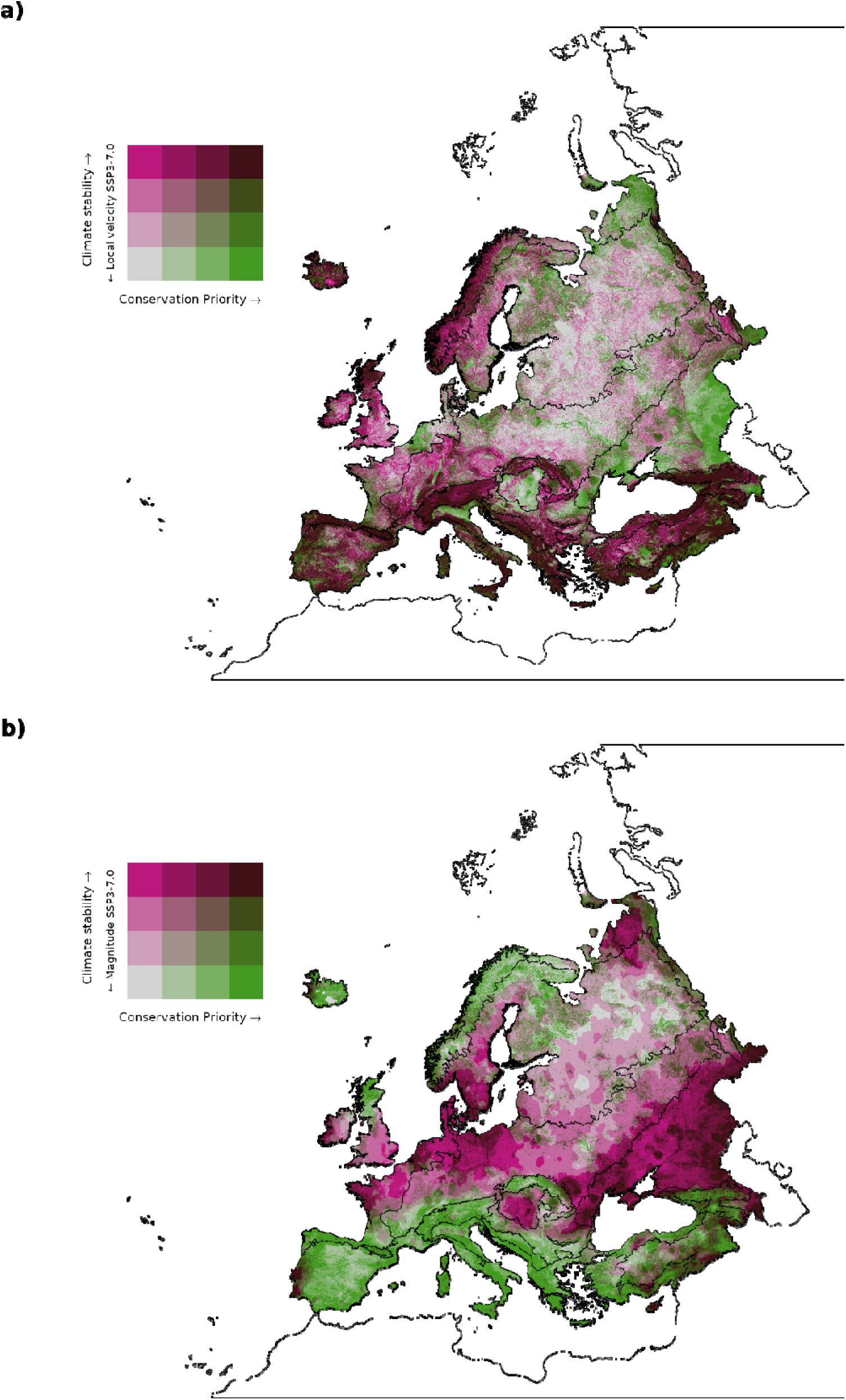
Maps of Europe illustrating the synergies (dark violet) and trade-offs (dark red and dark blue) between priorities to cover bioclimatic components and priority to maintain climatically stable areas under scenario SSP3-7.0. Panel a) represents climatic stability in terms of velocity; panel b) represents climate stability in terms of magnitude. In both panels, the horizontal axis of the legend represents conservation priority ranking (from grey to dark red), increasing from left to right, while the vertical axis represents climate stability, (from grey to blue) increasing from bottom (high values of local velocity or magnitude) to top (low values of local velocity or magnitude).

Trade-offs, instead, represent locations where either (i) high conservation priority overlaps with low climate stability (i.e., high velocity or magnitude), or (ii) low conservation priority overlaps with high climate stability. In the first case, areas of high conservation value and high climate velocity were mostly located in eastern Europe, along the borders of Russia and the Caspian sea, as well as in flat lowland regions of southern Europe, including the Po Valley in Italy, and large portions of Hungary and Romania (Fig. 4a). Areas with low priority but with high climate stability (low velocity) were predominantly in northern Europe, especially Scandinavia, the UK, Germany and northern France (Fig. 4a). When using climate magnitude as the climate stability metric, additional trade-off areas with high conservation values but high climate magnitude (low stability) were identified across the Mediterranean basin, Iceland, Scotland and northern Fennoscandia (Fig. 4b). Conversely, stable but low-priority regions, areas with low magnitude (high stability) but low conservation rank, were found across central Europe, stretching from east to west.

We further extracted two sets of priority areas, defined as the top 30% of areas in Europe with the highest prioritization value for bioclimatic components and simultaneously the lowest values of either climate velocity or magnitude (i.e. potential refugia). We found that a substantially larger number of components were underrepresented in these synergy areas (<10% species’ range in the final prioritization, Fig. S2.4) compared with selecting the same amount of land based solely on bioclimatic components. Specifically, selecting synergy areas based on climate velocity resulted in 595 components with low representation (Fig. S2.5), i.e. seven times more than in the original top-30% priorities areas (i.e. disregarding climate exposure). The effect was even larger when using climate magnitude, with 982 bioclimatic components underrepresented (i.e. 11 times more than under the original prioritisation). Coverage of different bioclimatic components was also markedly less equitable in the synergy-based prioritisations (Gini index = 0.312-0.450, when considering velocity and magnitude respectively; Fig. S2.6), compared to the biodiversity-only prioritization (Gini = 0.273).

To assess the spatial agreement between different conservation strategies, we quantified the relative overlap of areas selected under different prioritization approaches with the synergy areas. We found higher overlap between climatically stable areas and synergy areas (73.01% and 59.53% overlap for velocity and magnitude respectively), compared to biodiversity priority areas (67.26% and 48.38% overlap for velocity and magnitude respectively).

## Discussion

Our framework contributes to climate-resilient conservation by shifting the focus from static species distributions to the preservation of species bioclimatic niche diversity. By identifying and conserving distinct bioclimatic components within species’ ranges, we aim to safeguard the behavioural, phenotypical and genetic variation that underpins their adaptive capacity. This approach recognizes that species are not uniform units, but encompass populations adapted to different climatic conditions, an insight that becomes increasingly relevant as climate change accelerates ^15,23^.

The use of species-specific bioclimatic clustering allowed us to move beyond generalized biogeographic regions, or broad climate classifications. Instead of applying a uniform climate partitioning, we tailored the number of clusters to each species. Using self-organizing maps and k-means clustering, we identified biologically meaningful bioclimatic components ^28,29^. This flexible, species-centred approach allowed us to define an optimal number of clusters for each species, rather than applying a predefined scheme across all taxa ^15^. In doing so, we accounted for local adaptations and ecological specificity ^22^, capturing the range of climatic conditions within species’ distributions. This enables the representation of intra-specific climatic niche diversity—an essential component of evolutionary potential—by reflecting differences among local populations and their capacity to respond to environmental change.

The high-priority conservation areas identified through our framework are predominantly located in mountainous, coastal, and island regions, which are characterized by high climatic heterogeneity ^30,31^. The prevalence of high-priority areas in these regions likely stems from the presence of multiple narrowly ranged bioclimatic components, especially in mountainous zones where sharp climatic gradients and topographic complexity foster spatially restricted niches. Similarly, the edges of the European continent, such as the southern and eastern margins, may harbor unique bioclimatic components representing peripheral “tails” of broader Central Asian or North African climatic clusters, which are often spatially narrow and distinct, further driving their high prioritisation.

By integrating both intraspecific climatic variability and spatial patterns of climate stability, our approach addresses key uncertainties in conservation planning and contributes to the design of a climate-proof, ecologically coherent TEN-N. Importantly, we identified a number of synergy areas, regions where high priority for conserving bioclimatic niches overlap with high climatic stability. These “no-regret” zones are particularly valuable because they are expected to retain suitable environmental conditions over time while encompassing a wide range of bioclimatic components critical for species’ adaptive potential ^14,16^. Their alignment with current policy targets, such as the EU 2030 Biodiversity Strategy, makes them especially actionable for immediate protection. These findings align with growing calls to prioritize climate refugia in conservation planning ^32^ and directly support ongoing efforts to build a climate-resilient Trans-European Nature Network (TEN-N) ^1,33^.

While the protection of climate refugia is increasingly recognized as a cornerstone of climate-smart conservation ^13,34^, our results also caution that focusing on climatic stability alone might contrast with the objective of supporting species adaptation to climate change. In fact, some stable areas do not coincide with the most critical priority areas for representing species bioclimatic components. Relying exclusively on stable-climate refugia could therefore lead to underperforming efforts to conserve the full diversity and adaptive potential of species under climate change. Interestingly, synergies with local velocity provide more balanced representation of species components (and overall a lower number of poorly represented components) compared to synergies with climate magnitude (Fig. S2.5), indicating different trade-offs between biodiversity and climate refugia between locations that change slowly (low velocity) and those that change little (low magnitude). The results of the overlap of single strategy and synergy based solutions also highlight these trade-offs showing that velocity-based synergies align most with single-metric solutions, indicating strong spatial convergence between climatically stable and biodiversity-prioritized areas, while magnitude-based synergies show lower concordance, suggesting greater complementarity with biodiversity hotspots.

We found that pursuing climate stability comes at a particular cost of reduced representation for several biodiversity elements occurring in areas subject to substantial levels of climate exposure, i.e. “trade-off” areas. This means some species will require conservation measures in areas predicted to face substantial climate change. These areas pose complex challenges: while they harbour key bioclimatic diversity, their instability increases the risk of local extinctions ^9,35^. Nonetheless, they may also function as adaptation hotspots, where unique or vulnerable climatic niches could support evolutionary responses if appropriate measures are implemented. Dynamic conservation strategies will be required in these zones, including enhancing ecological connectivity ^36^, restoring degraded habitats, or, where necessary, considering assisted migration ^37^. Other trade-off areas include lowlands with high climate stability but low prioritization outcomes; these areas may hold strategic value under future climate scenarios, serving as colonization zones, or stepping-stones, for locally adapted genotypes. These trade-offs exemplify the limitations of static conservation approaches and underscore the importance of flexible, forward-looking planning to track shifting species distributions and climate trajectories ^13^.

The insights of our work call for more integrative conservation planning approaches capable of balancing climate resilience with biodiversity representation. Rigidly focusing on stable areas risks overlooking taxa that inhabit more exposed or dynamic environments— populations that may be crucial for evolutionary responses to climate change—areas where conservation interventions are most urgently needed. Despite their vulnerability, such areas can still support evolutionary processes and may harbour populations critical to long-term species adaptation ^38^. A promising path forward is a ‘bet-hedging’ strategy that combines the proactive protection of climatically stable refugia with reactive investment in high-risk areas that are critical for adaptation ^13^. Such a dual strategy can be further strengthened by establishing climate connectivity corridors, linking low-risk and high-risk zones to facilitate species’ range shifts and enhance resilience across fragmented landscapes.

## Conclusions

Our framework allows identifying synergy areas which maximise the representation of species-specific bioclimatic components while favouring areas of high climatic stability, to support climate-smart conservation planning. By explicitly incorporating intra-specific climatic niche diversity into spatial prioritization, the framework enhances our ability to identify areas that support both species persistence and adaptive potential under future climate conditions. This output can be integrated into existing policy instruments such as Natura 2000, national biodiversity strategies, and the Trans-European Nature Network (TEN-N)^33^, aligning directly with international conservation policy ^1,2^.

Practically, our findings caution against an over-reliance on climate stability alone. While refugia are vital for long-term persistence, they do not capture the full breadth of biodiversity or adaptive potential that conservation must safeguard. Effective strategies will require a balanced approach, combining proactive protection in stable climate areas with dynamic, responsive actions in exposed areas. By offering an evidence-based means to integrate current priorities with future resilience, the framework supports a more strategic, adaptive, and future-oriented vision for biodiversity conservation in Europe and beyond.

## Methods

### Species distribution models

We modelled the distributions of 1,207 terrestrial vertebrate species (117 amphibians, 526 birds, 296 mammals and 268 reptiles) across Europe and adjacent regions using environmental predictors describing climate (CHELSA) ^39^, topography (EarthEnv; Amatulli et al. 2018), soils (SoilGrids; Poggio et al. 2021), hydrography ( EU-Hydro, HydroSHEDS, GLWD) ^42–44^, land cover (CORINE, 2020,ESA CCI 2016) ^45,46^, and land use ^47^ at 1 km² resolution. Species occurrences in the period 1980-2023 were obtained from GBIF, filtered for potential errors using CoordinateCleaner ^48^, and validated against ranges from IUCN and BirdLife International. Records outside expert ranges were excluded, with small buffers added to account for uncertainty in range assessments. Pseudo-absences were sampled within and outside these buffered ranges, weighted by target group sampling effort, and repeated to reduce selection bias. Species distribution models were fitted using Random Forest ^49^, XGBoost ^50^, and neural networks ^51^. Data were partitioned into five spatial blocks for cross-validation ^52^. Up to 75 models per species (3 algorithms × 5 pseudo-absence sets × 5 folds) were trained, and only those with a True Skill Statistic (TSS) > 0.4 ^53^ were retained. Ensemble suitability maps were then computed as the mean prediction across retained models, with values ranging from 0 (unsuitable) to 1 (highly suitable). To approximate realized ranges, suitability maps were constrained by expert ranges smoothed with an exponential decay kernel, and converted into binary predictions using species-specific thresholds optimised with the Continuous Boyce Index ^54^. The latter identifies the suitability value above which presences are consistently more frequent than expected under a null model.

### Defining species bioclimatic components

We identified individual bioclimatic components within each species distribution based on a cluster analysis. Our purpose was to represent the diversity of climatic conditions that each species occupies, in order to represent the variability in their realised climatic niches ^15,23^. This variability is expected to capture species regional ecological adaptations to climatic conditions, which can be manifested through: morphological differences ^55–57^, dietary differences ^58–60^ or behavioural differences ^61–64^. Capturing these differences ensures that the evolutionary adaptive capacity of a species to climate change is preserved. We did not apply the framework on 157 species with restricted geographic ranges (range size smaller than 20,000 km²)^27^, to prevent excessive fragmentation of their conservation priorities.

For the remaining 1,050 species, we extracted the values of relevant bioclimatic variables ^39,65^ across the predicted distribution range. We selected the same bioclimatic variables as in the SDMs, specifically: temperature seasonality (bio4), annual precipitation (bio12), precipitation seasonality (coefficient of variation, bio15), growing degree days heat sum above 5°C (gdd5), snow cover days (scd) and snow water equivalent (swe). To broaden the framework’s applicability, when SDMs are unavailable, we recommend selecting bioclimatic variables that capture key climatic dimensions—thermal conditions, water availability, and critical seasonal periods—relevant to the species’ ecological and physiological responses to climate change. After extracting the bioclimatic variable values across all pixels of species predicted distributions, we employed Self-Organizing Maps (SOMs) to identify a reduced set of local climate nodes that represented the full spectrum of climatic conditions of the species. To reduce the dimensionality of the environmental data while retaining its complexity, we applied Self-Organizing Maps (SOMs), a form of unsupervised artificial neural networks that identify local climate nodes across species distributions ^66,67^. SOMs project high-dimensional input data—here, the bioclimatic values across species’ ranges—onto a two-dimensional grid where nodes (or neurons) represent typical combinations of bioclimatic conditions. The weight vectors of the nodes are iteratively adjusted to reflect the structure of the data, resulting in a spatial organization where similar conditions are mapped to nearby nodes ^28,29,68^. The number and layout of SOM nodes for each species were determined using heuristic ^69–71^ based on the number of cells in the species’ range ^69–71^, with a maximum of 1 million randomly sampled cells used for species with very large ranges. All SOMs were implemented in R using the Rsomoclu package ^72^.

We then applied k-means clustering to the SOM nodes to delineate distinct bioclimatic components for each species ^28^. The optimal number of clusters was determined using four complementary validation indices—silhouette coefficient ^73^, Dunn index ^74^, Davies–Bouldin index ^75^, and Calinski–Harabasz index ^74^—implemented through the R package NbClust ^76^. These metrics assess clustering quality by evaluating intra-cluster compactness and inter-cluster separation, and we used the best values across indices to derive a consensus estimate of the optimal number of clusters per species. A minimum of three clusters was enforced to avoid oversimplification ^77^. Thus, in order to increase the interpretability of clustering results we used the *FeatureImp* function to find the variables which drive the cluster assignment and score them according to their relevance ^78^. Specifically it computes the permutation missclassification rate for each variable of the data, then the mean misclassification rate over all iterations is interpreted as variable importance.

Once the optimal number of clusters was defined, species distributions were spatially classified into bioclimatic components. To do this, we trained a K-Nearest Neighbours (KNN) model to assign each grid cell to the appropriate cluster based on the SOM output. The SOM nodes were split into training (70%) and testing (30%) subsets, and we used the caret package in R to tune the number of neighbors (k) via 10-fold cross-validation, selecting the best model using accuracy and Kappa statistics. The final KNN model was then used to classify all grid cells across each species’ distribution, resulting in a set of raster maps that delineate bioclimatic components, which were subsequently used in the spatial prioritization analysis.

### Climate risk

Climate exposure, and conversely, refugia, can be identified using various climate-based metrics ^12,34^ most commonly climate velocity and climate magnitude ^12,38,79,80^.

We used two climate exposure metrics ^12^ to assess climate risks under scenario SSP3-7.0: local climate velocity (sensu Loarie et al., 2009) ^35^, and the magnitude of change (sensu Williams et al., 2007) ^79^. Usually, multiple climate change scenarios are recommended to capture the range of possible future conditions and associated uncertainties ^81,82^, but SSP3-7.0, as a high-emission, high-uncertainty pathway marked by regional fragmentation, weak climate policies, and major land-use changes, is particularly relevant for biodiversity and ecosystem assessments under a relatively realistic future scenario.

Local climate velocity represents the speed and direction at which species must move to remain within suitable climatic conditions, describing the intensity of change in a given location relative to local variation in climatic conditions, providing insights into species’ potential to relocate ^35^. We calculated the local velocity, following the gradient-based approach ^83^ and using the R package *VoCC* version 1.0.0 ^83^.

The second metric, climate change magnitude describes how much the climatic conditions of a location (focal cell) change over time, standardized by historical interannual variability ^84^. It reflects the degree of climate change exposure for species at a given site ^79,85^, and is quantified using the standardized Euclidean distance (SED) between baseline (1981–2010) and future (2041–2070) climates ^79,86^, (Appendix S1).

### Spatial prioritisation

We used Zonation 5 software v2.1 to identify priority areas for the conservation of bioclimatic components across Europe. This software’s priority ranking method has been shown to effectively achieve comprehensive biodiversity coverage, including for narrow-range features ^87^, which is important for capturing ecologically unique bioclimatic components. Zonation ranks grid cells based on their conservation importance by evaluating each input feature’s (here, bioclimatic components) representation within the grid cell, the complementarity between cells, and applying an iterative conditional sort algorithm ^88^. The ranking process starts with an initial arrangement of grid cells, which is then refined iteratively according to the marginal loss rule, which defines how feature values are combined into a single, cell-specific numerical value. This allows the algorithm to compare, rank, and order grid cells from least to most important (see Moilanen et al. 2022 for methodological details). Once the process stabilizes, Zonation produces a priority ranking for each pixel, highlighting areas with the highest relative representation values for the bioclimatic components. We used the ‘CAZ2′ as the marginal loss rule option, which results in both relatively high mean coverage and high minimum feature representation in the top-ranked areas ^88^. This option supported our goal of identifying areas that are both ecologically diverse (i.e., high average representation of all bioclimatic components) and balanced in feature representation (i.e., minimizing the loss of the least-represented bioclimatic components). For the purpose of our analysis we select the top-30% priority areas, based on the 30×30 target ^2^.

### Measuring the equitability of components coverage

To assess how equitably conservation prioritization represents the coverage of each bioclimatic component, we calculated the Gini index, a standard measure of inequality ranging from 0 (perfect equality—all components equally represented) to 1 (maximum inequality—only one component is represented). In this context, lower Gini values indicate more even representation of a species’ bioclimatic diversity, while higher values reflect spatial bias, i.e. overrepresentation of some components and neglect of others.

For each prioritization strategy – biodiversity components and biodiversity-climate synergies – we extracted the equitability in the representation of each component computing the Gini coefficient using the *Gini* function in R ^89^.

We also plotted Lorenz curves for each strategy at a 30% coverage level (0.7 priority rank) to visually examine the cumulative distribution of subcomponent representation. The Lorenz curve is a graphical representation used to illustrate the degree of inequality within a distribution. It plots the cumulative proportion of a variable (e.g., coverage of bioclimatic components) against the cumulative proportion of the total number of elements (e.g., all components), ordered from the least to the most represented. A perfectly equal distribution would follow a straight diagonal line (the line of equality), indicating that all elements contribute equally. Deviations below this diagonal indicate increasing inequality — the further the Lorenz curve bows away from the line of equality, the more uneven the distribution. In this study, the Lorenz curve is used to visualize how evenly each conservation strategy represents the climatic components of species: curves closer to the diagonal indicate more balanced representation, whereas more curved lines reveal a stronger bias toward certain components.

### Measuring spatial overlap between strategies

We quantified to what extent the synergy-based prioritizations captured areas also prioritized under single-strategy approaches. Specifically, we compared (i) the biodiversity-only prioritization with the synergy solution incorporating climatic variability and stability (biodiversity–velocity synergy, biodiversity–magnitude synergy), and (ii) the climate-only prioritization (low velocity or low magnitude) with their corresponding synergy solution. For each pairwise comparison, binary maps of selected priority areas (based on equivalent target levels) were intersected, and the proportion of spatial overlap was calculated as the area shared between the two strategies divided by the total area of the reference strategy.

## Acknowledgements

This research was funded from the project NaturaConnect, which receives funding under the European Union’s Horizon Europe research and innovation programme under grant agreement number 101060429. Views and opinions expressed are however those of the authors only and do not necessarily reflect those of the European Union or the European Research Executive Agency. Neither the European Union nor the granting authority can be held responsible for them. The authors wish to acknowledge CSC – IT Center for Science, Finland, for generous computational resources.

