## Supplementary Material for "Synergies and trade-offs between maintaining climate niche variability and preserving climate stability"

### **Appendix S1.**

### **Supplementary Methods**

We proposed a framework to define spatial priorities for the conservation of species individual bioclimatic components, which we then contrasted with estimates of climate stability in Europe (Fig. 1, main text). We started from the distribution map of species, here derived from species distribution models (SDMs), and identified climatically distinct clusters within each species' distribution which we refer to as bioclimatic components. We then run a spatial prioritization analysis to identify priority areas for the conservation of tetrapod niches across Europe.

We contrasted these areas with maps representing climate stability (in terms of low velocity and low magnitude) to identify synergies and trade-offs between the conservation of areas that maintain climate niche variability for species and the conservation of areas with high climate stability.

### **Defining species bioclimatic components**

We identified individual bioclimatic components within each species distribution based on a cluster analysis. Our purpose was to represent the diversity of bioclimatic conditions that each species occupies, in order to represent the variability in their realised climatic niches ^1^. This variability is expected to capture species regional ecological adaptations to climatic conditions, which can be manifested through: morphological differences ^2–4^, dietary differences ^5–7^, behavioural differences ^8–11^. Capturing these differences ensures that the evolutionary adaptive capacity of a species to climate change is preserved. We did not apply the framework on 157 species with restricted geographic ranges (range size smaller than 20,000 km²)^12^, to prevent excessive fragmentation of their conservation priorities.

For the remaining 1,050 species, we extracted the values of relevant bioclimatic variables ^13,14^ across the predicted distribution range. We selected the same bioclimatic variables as in the SDMs, specifically: temperature seasonality (bio4), annual precipitation (bio12), precipitation seasonality (coefficient of variation, bio15), growing degree days heat sum above 5°C (gdd5), snow cover days (scd), snow water equivalent (swe). To broaden the framework's applicability, when SDMs are unavailable, we recommend selecting bioclimatic variables that capture key climatic dimensions—thermal conditions, water availability, and critical seasonal periods—relevant to the species’ ecological and physiological responses to climate change.

After extracting the bioclimatic variable values across all pixels of species predicted distributions, we employed Self-Organizing Maps (SOMs) to identify a reduced set of local climate nodes that represented the full spectrum of climatic conditions of the species. This step was necessary for dimension reduction before the clustering analysis (see below). SOMs are a form of artificial neural networks that utilize unsupervised learning and adaptive weights for clustering data into local nodes ^15^. This approach has minimal reliance on parameter settings and does not require predefined similarity measures for grouping data. It is also capable of detecting isolated patterns and structures within the data ^16^. Specifically, SOMs map multidimensional data onto a two-dimensional X-Y plane, known as the SOM grid, which consists of nodes or neurons. These nodes classify locations with similar bioclimatic conditions ^17,18^. Each node is defined by a model weight vector, representing the centroid means in the SOM solution or the distance of the node from the input data—in this case, the values of bioclimatic variables within species' ranges. During the learning process, the model's weight vectors are gradually adjusted to match the samples, ensuring they accurately reflect the underlying properties of the data. As a result, input vectors that are closer in the input space are mapped to nodes that are similarly close on the grid map ^19^. The nodes are then arranged on the grid such that more similar ones are positioned near each other ^20^.

The first step to run a SOM is to determine the number of map units, which should be set as large as possible to preserve mapping smoothness and enhance generalization while being computationally tractable. To determine the number of map units for each species, we applied an heuristic rule ^21^ suggesting a number of nodes that is five times the square root of the number of samples, or in our case, number of grid cells that falls inside the species SDM ^22,23^. Next, the map dimension ratio, the ratio of the grid side lengths representing the node mapping (i.e., how many units are along each axis of the SOM grid) is determined by the ratio between the two largest eigenvalues of the training data ^20^. Due to computational constraints, we randomly sampled 1 million grid cells from the distributions of large-ranged species (distribution >1 million km²) to run SOM, using the entire distribution of all other species (distribution <1 million km^2^). SOM analyses were produced using the R package *Rsomoclu* version 1.7.6 ^24^.

The second step involved applying k-means clustering to the local climate nodes generated by the SOM ^20^. The optimal number of clusters for k-means was determined using the R package “NbClust” version 3.0.1 ^25^, considering four different cluster validation indices: silhouette coefficient (SC), Dunn index (DI), Davies–Bouldin index (DB), and Calinski–Harabasz index. The silhouette coefficient evaluates clustering quality by combining intra-cluster cohesion and inter-cluster separation. It measures how similar an object is to its own cluster compared to other clusters, with values ranging from -1 (indicating potential misclassification) to 1 (indicating well-placed observations); values around 0 suggest that an observation lies between two clusters ^26^.

The Dunn index identifies clusters that are both compact and well-separated, calculated as the ratio between the minimum inter-cluster distance and the maximum intra-cluster distance, with higher values indicating better clustering ^27^. The Calinski–Harabasz index measures the ratio of between-cluster to within-cluster dispersion, with higher values denoting more clearly defined clusters meaning that data points within each cluster are close to each other, and the different clusters are well-separated from each other ^28^. The Davies–Bouldin index quantifies the similarity between clusters by assessing the ratio of within-cluster scatter to between-cluster separation. Lower values indicate better clustering results, with values near 0 reflecting well-separated and compact clusters ^29^, but in this context, a value around 1 still suggests that the clusters are fairly separated and compact.

Since these indices evaluate different and complementary aspects of clustering performance—such as compactness, separation, and overall cluster definition—we computed the optimal number of clusters for each species separately based on the best value of each index. We then averaged the resulting values to derive a final, consensus estimate of the optimal number of clusters for each species.

After selecting the optimal number of clusters per species and performing the k-means clustering, we assessed the quality or validity of the resulting clusters by recalculating the aforementioned metrics. To prevent oversimplification of species' bioclimatic components, we set a minimum of three clusters ^30^.

The result is a minimal set of clusters that represent the primary bioclimatic components within species distributions. After identifying the optimal number of bioclimatic clusters, the current species distributions were divided according to this, producing bioclimatic component-specific maps for each species.

To divide the distributions, we used K- Nearest Neighbours (KNN) to reclassify the entire current species distributions by assigning each pixel to a cluster, based on the labels of the closest neighbours in the training dataset (SOM nodes). Specifically, KNN determines the classification or grouping of an individual data point based on its proximity to others. The classification is based on a majority vote from its neighbours, with the data point being assigned to the most common class among its 'k' nearest neighbours, where 'k' represents the number of neighbours considered in the analysis. Thus, we split resulting SOM nodes into training (70%) and testing (30%) subsets, and used the caret package to train a KNN model using a 10-fold cross-validation. We tuned the number of neighbours (k) by testing values from 1 to 10 and then evaluated the model's performance on a test set calculating accuracy and Kappa statistics. Thus, we used the final KNN model to assign each pixel from a species distribution into a bioclimatic cluster, finally producing a number of separate rasters representing bioclimatic components for each species to be used in the subsequent spatial prioritization analysis.

We also evaluated the importance of individual features in the clustering by measuring how cluster assignments change when each feature is randomly shuffled, using the *FeatureImp* function from the “FeatureImpCluster” R package ^31^. The function first establishes baseline clusters, then iteratively permutes each feature and re-clusters the data. It calculates the misclassification rate—the proportion of observations that switch clusters compared to the baseline. After all iterations, the function returns misclassification rates for each feature: higher values indicate greater importance, while lower values suggest minimal impact. This provides a quantitative assessment of which features most influence the clustering outcome.

### **Climatic metric calculation**

We measured the local velocity of change following the gradient-based approach ^32^, and using an automated workflow based on the R functions *tempTrend* and *spatGrad* in the “gVoCC” package ^33^. The velocity of change is defined as the ratio between the temporal trend of a climate variable, the rate of change of the variable through time, for baseline (1980-201) and future (2041 – 2070) periods (estimated as a regression slope), and the corresponding local spatial gradient of that variable (i.e., the vector sum of longitudinal and latitudinal pairwise differences at each focal cell using a 3 x 3-cell neighbourhood), described in Eq. [1].

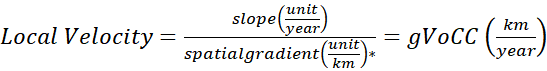
 [1]

**unit/year*: (°C/km for temperature and mm/km for precipitation)

The second metric, climate change magnitude, measures change in a climate parameter over time at a given location, providing information on the extent of the change in each location in relative terms of the climate variables as they are standardized by historical interannual climate variability ^34^. Ecologically, this metric provides direct information on the climate change to which a species is exposed in a given location ^35,36^. We quantified magnitude as dissimilarities between baseline (1981−2010) and future (2041–2070) climate by using the standardized Euclidean distance (SED) ^35,37^, as in Eq. [2].

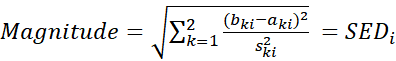
 [2]

For further details on methodological approach for the calculation of climatic metrics see Cimatti et al. 2025 ^38^.

### **Appendix S2.**

### **Supporting Results**

### **Identification of bioclimatic range components**

The median silhouette value was S = 0.32 (sd=0.054), with a 3.31% negative value (sd=2.03%), indicating that a very small number of grid cells within species distribution had an uncertain classification (e.g. between two possible clusters). This result indicates good overall coherence of the distribution grid cells within each cluster (Figure S2. 1b, 2c). The mean Dunn index value was D = 0.008, which indicated weakly compact and separated clusters, while Calinski Harabasz was relatively high (2191.28), suggesting that the proposed clusters were well-defined. The Davies–Bouldin index had a mean value of 1.19 which indicates a fair clustering quality.

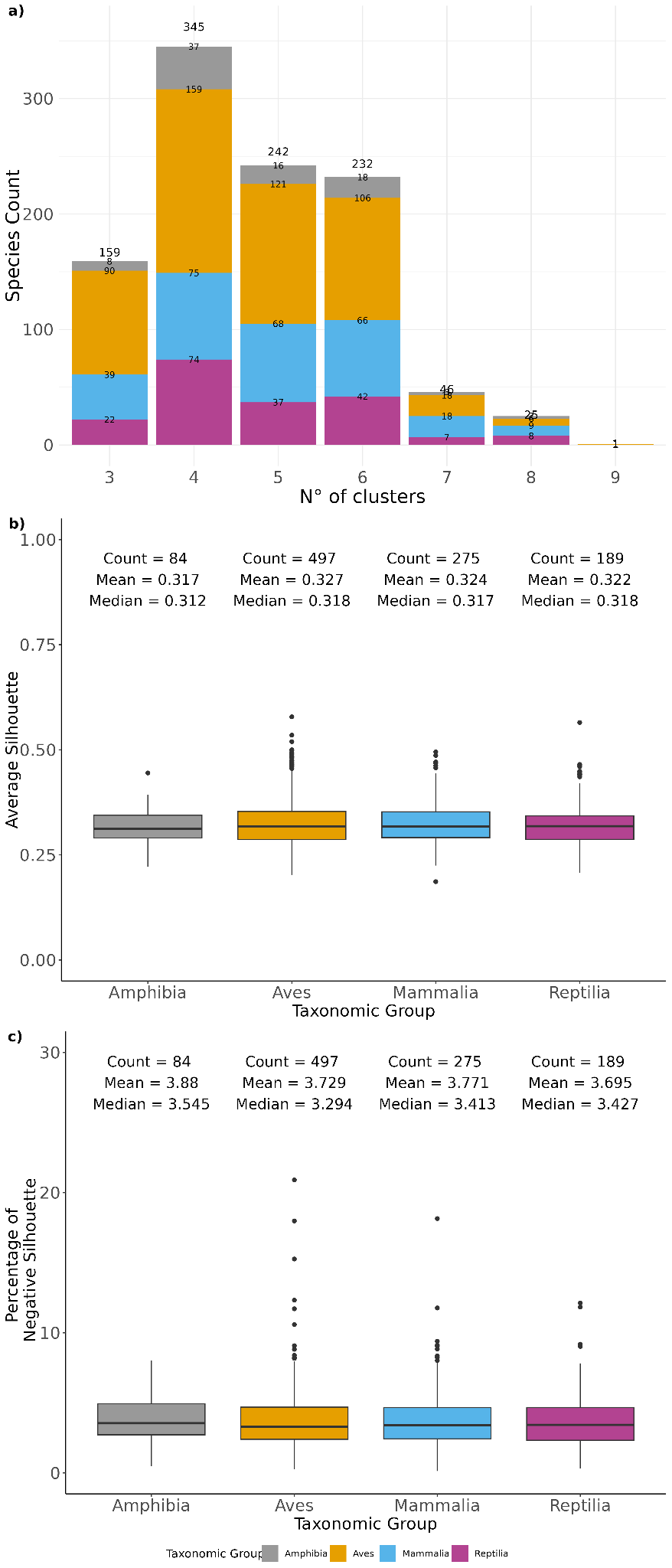

***Fig. S2.1:*** *Niche splitting results – a) Barplot showing the frequency of bioclimatic components (clusters) across species colored by the main taxonomic group. b) Boxplot showing clustering performance, reporting mean silhouette index value per species grouped and colored by taxonomic group. c) Boxplot showing the percentage of negative silhouette value per taxonomic group*.

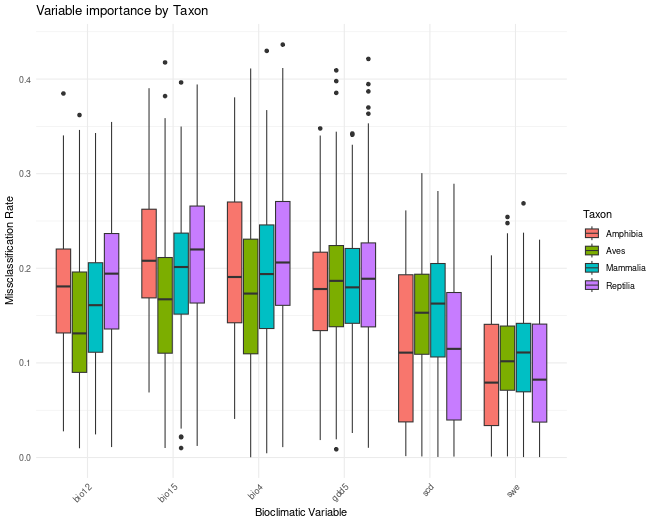

***Fig. S2.2****: Variable importance analysis across taxonomic groups (Amphibia, Aves, Mammalia, Reptilia) showing misclassification rates for key bioclimatic variables. Higher values indicate greater influence of each variable in defining ecological clusters for the respective taxa.*

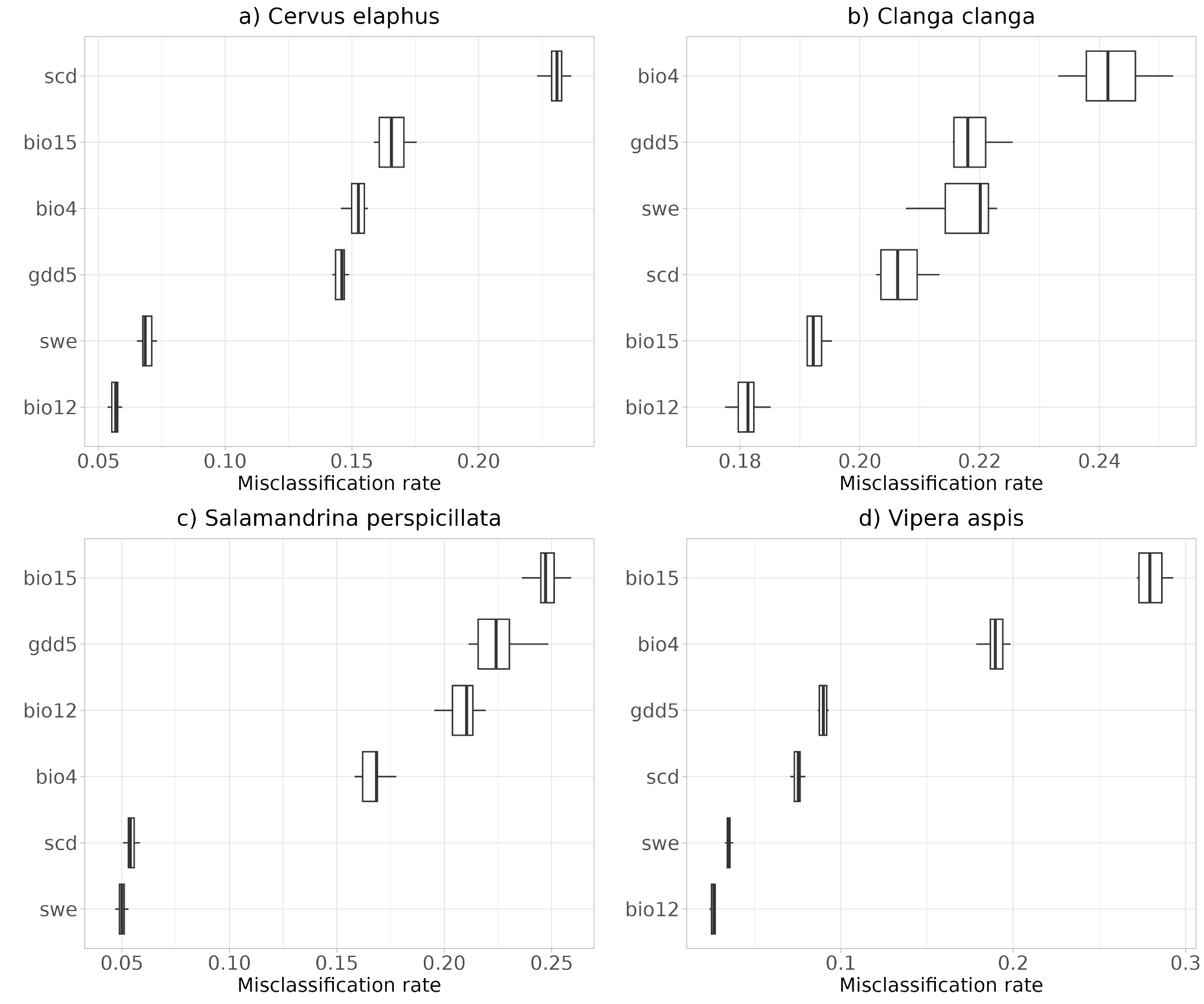

***Fig. S2.3****: Feature importance analysis across four species a) Cervus elaphus, b) Clanga clanga, c) Salamandrina perspicillata, and d) Vipera aspis) showing misclassification rates for key ecological variables.*

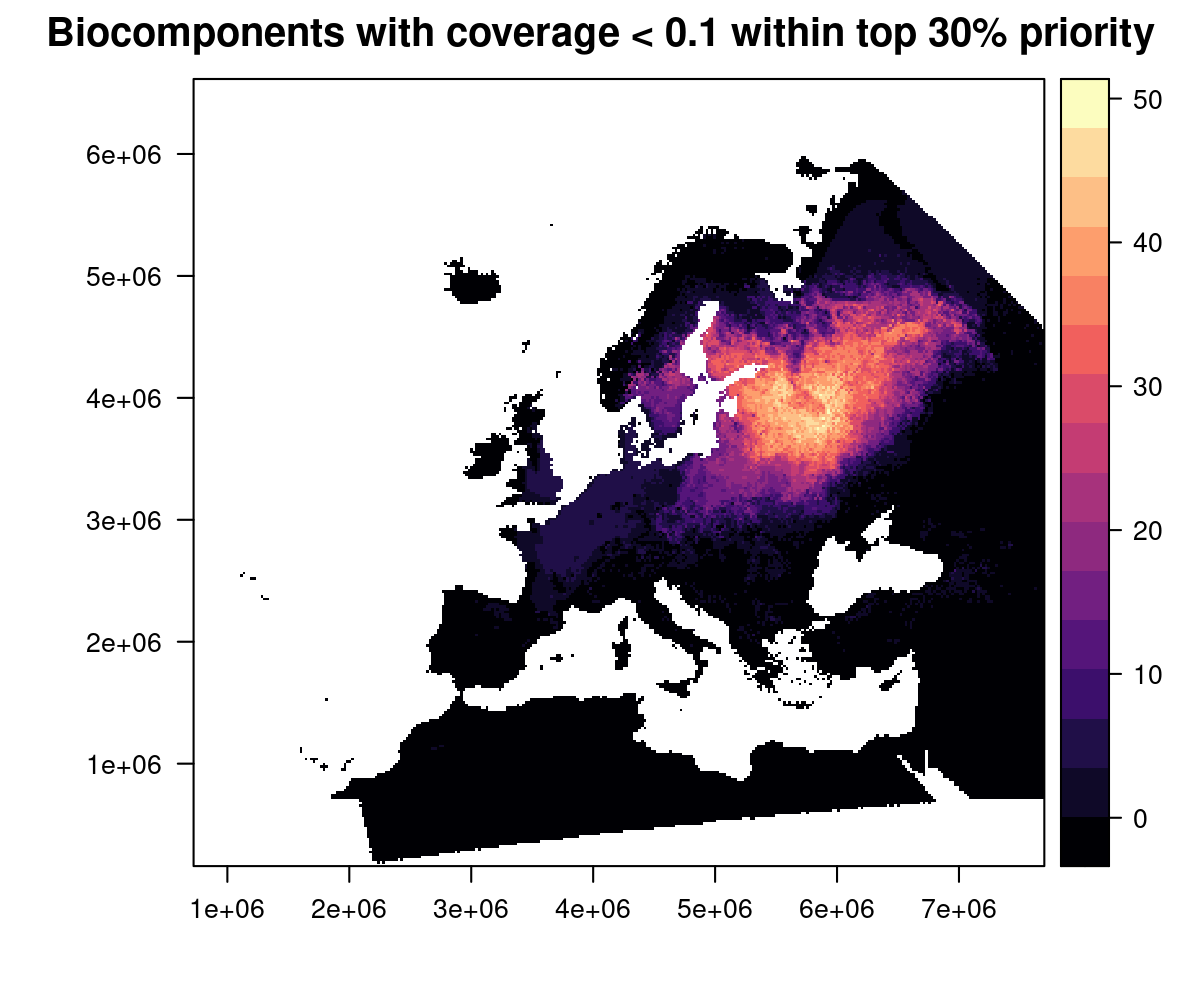

### ***Fig. S2.4:*** *Spatial distribution of bioclimatic components with low representation (coverage < 0.1) within the top 30% priority rank across Europe. Warmer colors indicate regions where a higher number of poorly represented biocomponents overlap.*

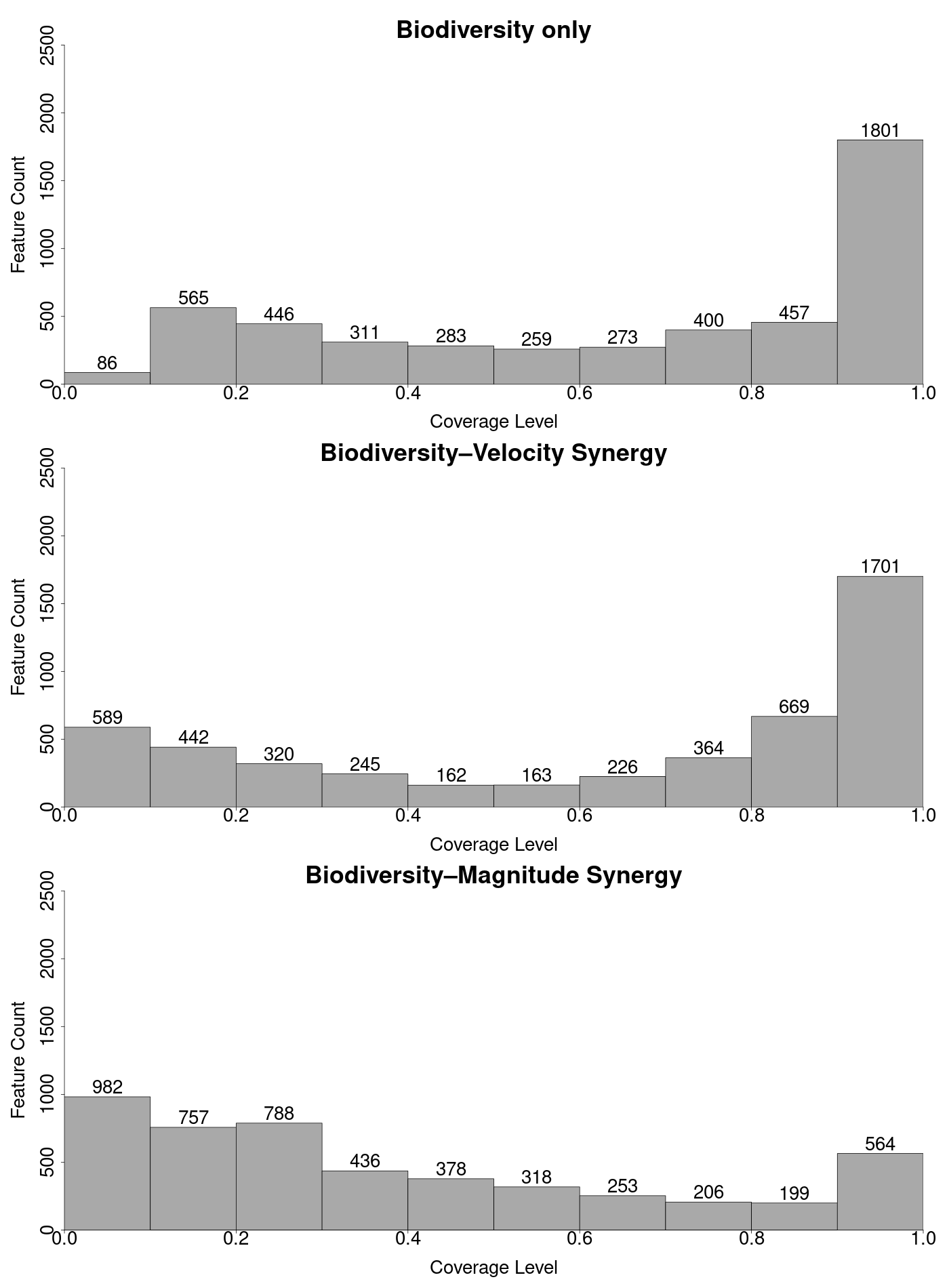

***Fig. S2.5:*** *Comparison of species coverage between biodiversity-only and combined biodiversity-climate stability prioritization scenarios (biodiversity–velocity synergy, biodiversity–magnitude synergy. The panel shows feature counts (species) at varying coverage levels (0.0–1.0) for different runs of zonation analysis.*

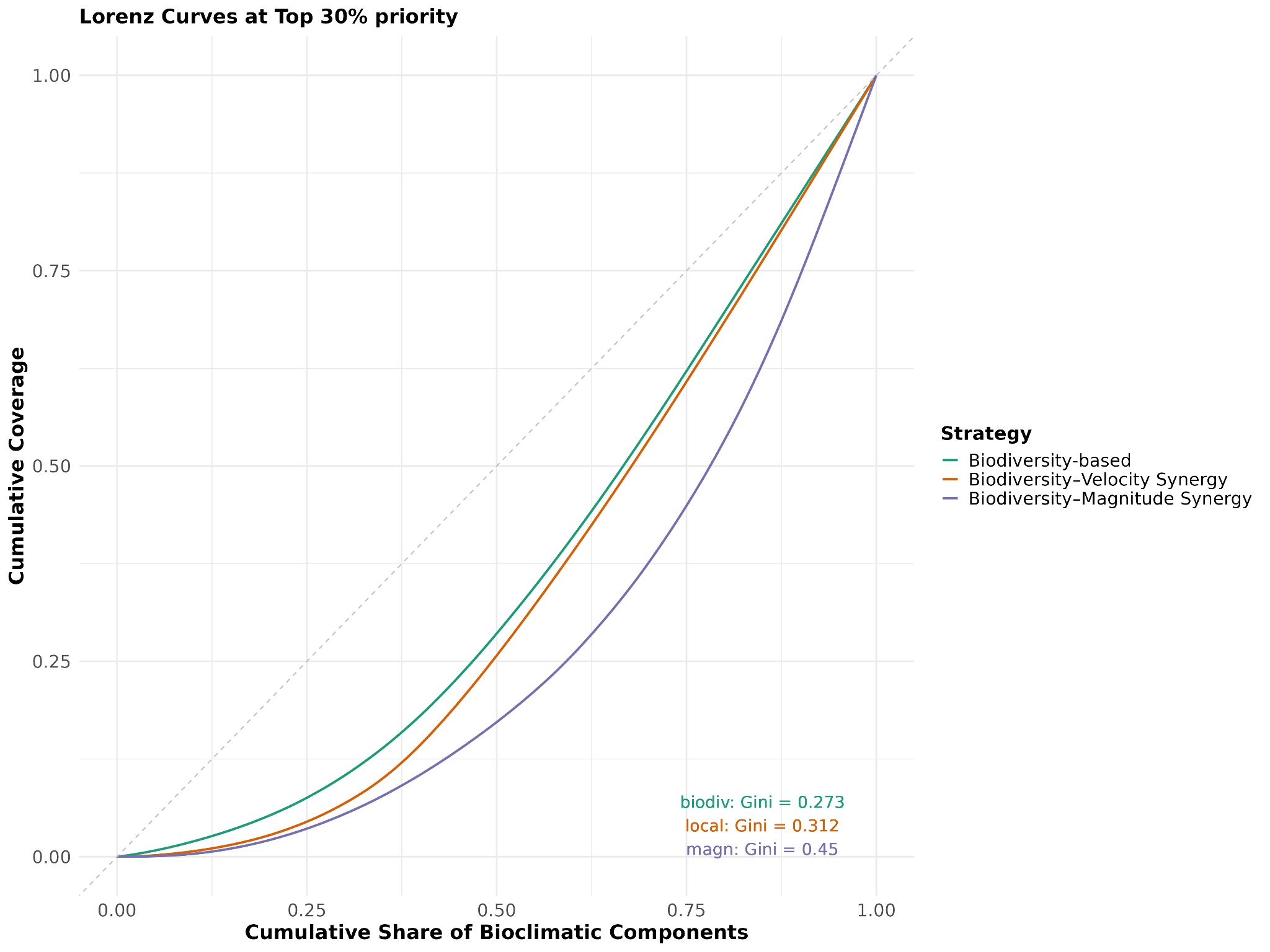

***Fig. S2.6:*** *Equity and distribution of conservation priorities under alternative strategies. Lorenz curves show the cumulative proportion of bioclimatic component coverage as a function of the cumulative proportion of area protected for three strategies (biodiversity, biodiversity–velocity synergy, and biodiversity–magnitude synergy) at the top 30% target Higher Gini values indicate higher inequality in the coverage of bioclimatic components.*
